# Complexity in SARS-CoV-2 genome data: Price theory of mutant isolates

**DOI:** 10.1101/2020.05.04.077511

**Authors:** Saurav Mandal, R.K. Sanayaima Singh, Saurabh Kumar Sharma, Md. Zubbair Malik, R.K. Brojen Singh

**Affiliations:** School of Computational and Integrative Sciences, Jawaharlal Nehru University, New Delhi-110067, India; School of Computer and Systems Sciences, Jawaharlal Nehru University, New Delhi-110067, India

**Keywords:** SARS-CoV-2, Multifractal analysis, Hurst exponent, Generalized dimension, Price equation

## Abstract

SARS-CoV-2 is a highly virulent and deadly RNA virus causing the Covid-19 pandemic and several deaths across the world. The pandemic is so fast that any concrete theory of sudden widespread of this disease is still not known. In this work, we studied and analyzed a large number of publicly available SARS-CoV-2 genomes across the world using the multifractal approach. The mutation events in the isolates obey the Markov process and exhibit very high mutational rates, which occur in six specific genes and highest in *orf1ab* gene, leading to virulent nature. *f* (*α*) analysis indicated that the isolates are highly asymmetric (left-skewed), revealing the richness of complexity and dominance by large fluctuations in genome structure organization. The values of *H_q_* and *D_q_* are found to be significantly large, showing heterogeneous genome structure self-organization, strong positive correlation in organizing the isolates, and quite sensitive to fluctuations in and around it. We then present multiple-isolates hosts-virus interaction models, and derived Price equation for the model. The phase plane analysis of the model showed *asymptotic stability* type bifurcation. The competition among the mutant isolates drives the trade-off of the dominant mutant isolates, otherwise confined to the present hosts.

## Introduction

The special concern about the Covid-19 is the type of pneumonia, which is deadly and highly infectious [1] causing the sudden outbreak of this disease from Wuhan, Hubei Province, China [1, 2] and, spread to different parts of the world that led to WHO to declare this epidemic as an emergency in global public health [3]. Coronaviruses (CoVs) belongs to *Coronaviridae* subfamily (which have *α, β, γ, δ*-CoVs genera) in the family *Coronaviridae* of the *Nidovirales* order [4]. The Covid-19 causing virus *SARS-CoV-2* is the seventh member in *β*-CoV genus [5], which can infect humans [6], and it’ s genetic manipulation studies, and sequence analysis has led to the claim that it is not laboratory manipulated virus [6, 7]. It’ s genome is +ssRNS (single-stranded positive-sense RNA virus) [4, 5] having 30kb genome size [8] which becomes 100-160nm in compact form [4, 6, 7]. Further, it’ s genome has been found 80% similarity with that of SARS-CoV, whereas around 96% similarity with that of bat coronavirus [9].

Clinical symptoms of Covid-19 include dry cough, sore throat, headache, fever, problems in breathing, dyspnea [3], which may lead to developing problems of organ failure, severe pneumonia, septic shock, acute respiratory distress syndrome, etc. to some patients [10]. However, SARS-CoV-2 enter and infect target host cells (lung, intestine, kidney, and blood vessels) through the receptor angiotensin-converting enzyme 2 (ACE2), and ACE2 expression is found to be increased in diabetes (type 1 or type 2) [11]. Further, hypertension is treated with ACE inhibitors and angiotensin II type-I receptor blockers (ARBs) [11], resulting in an increase in ACE2 expression [12]. Hence, diabetes and hypertension patients might have a severe risk of developing Covid-19 as these patients use ACE inhibitors and ARBs as a drug which upregulate ACE2 expression [13]. Moreover, there are number of reports and unsolved cases regarding the infection of this virus to different people at different geographical conditions, people of different ages, sex and pregnant woman, demographical parameters, etc. [3, 14, 15].

There is no concrete theory so far why SARS-CoV-2 has a high affinity to human host cells, which in turn led to extremely fast human-to-human transmission as compared to other members of CoVs and even among seven members in β-CoV. One belief is that SARS-CoV-2 has receptor binding protein (RBD) that has high cohesive towards human receptor ACE2 [7, 9], it’ s spike protein has functional polybasic cleavage site at the S1-S2 junction which is quite variable [7, 9, 16]. Then it might allow jumping to the human host cells via comfortable adap-2tation for fast transmission [7]. However, theory of the origin of this virus and pandemic caused by it are still debatable.

### Winsor theory of Covid-19 pandemic

The initial growth of the infected people of Covid-19 pandemic in any country across the world follow interesting Malthusian exponential growth law with time [17] as shown in Fig. 1 adapted from Roser et. al. world data [18]). If *N*(*t*) is the time-dependent confirmed SARS-CoV-2 infected population (Fig. 1A), or confirmed death population (Fig. 1B), one can see that each curve of each country follow one of the following growth law at different growth stages [17, 19–21],

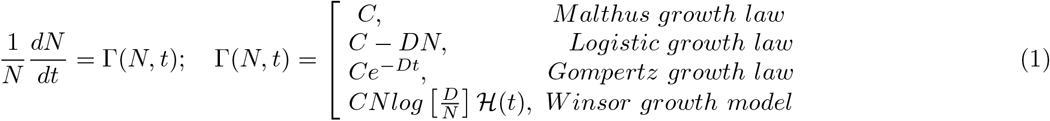

where, *C* and *D* are constants, and 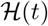 is a function of *t*. The initial growth rate of Covid-19 pandemic is dominated by *Malthus growth rate* [17], and if we consider significant amount of time of the pandemic, coronavirus infected or death population nicely follow *logistic* [19] or more systematically *Gompertz growth law* [20]. The spread of every disease epidemic or pandemic generally increase monotonically (following Malthus law [17]) due to availability of large susceptible population till it reach threshold value [22] (follow logistic or Gompertz law [19, 20]). After reaching this threshold size of epidemic, it will gradually decrease the size of epidemic due to removal of all susceptible populations and gradual decrease of the virus virulence, die out slowly [22]. In such scenario, the Winsor growth model [21] is quite suitable to explain and forecast, because, the function 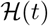 in the solution of Winsor model in equation (1), 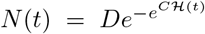, can have the change in sign, and can be expanded through Taylor series, 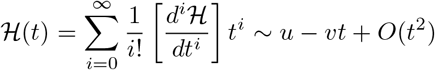 (one can see the change in sign in the second term). Thus, we have Winsor growth law, *N*(*t*) = *De^-e^u-vt^^*, which is modified form of Gompertz model, can well explain the pandemic. The uncontrolled fast growth of *N*(*t*) due to lack of proper checking in the *N*(*t*) outgrowth may cause inevitable shortage of food, trapping in (starvation-*N*(*t*) outgrowth) cycle which may lead to fast decrease in *N*(*t*), the condition known as *Malthusian catastrophe* given by, 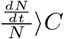 [17, 23]. However, happening of this condition is unlikely, even though the dynamical process could lead to the sacrifice of a certain population of infected people *N_U_*(*t*), who is most probably unhealthy/non-immune population out of *N*(*t*), and the most fitted one *N_F_*(*t*) = *N*(*t*) — *N_U_*(*t*) will survive. But, the question of why the growth rates of both infected and death are different in different countries at the epidemic network level or genetic level and how to control the epidemic/pandemic size are still open questions.

**FIG. 1:**
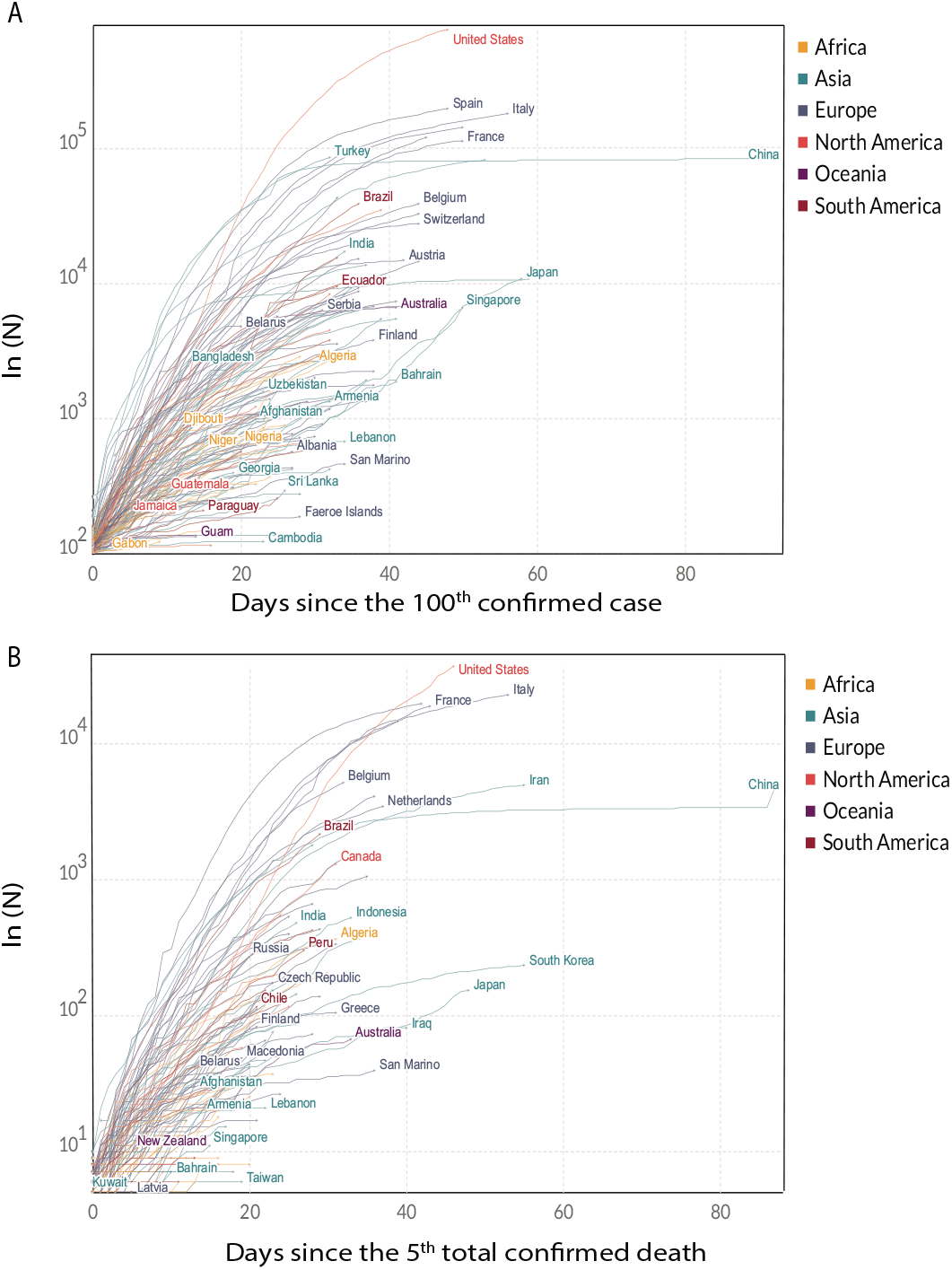
Dynamics of Covid-19 pandemic: The dynamics of country wise confirmed (A) infected and (B) death cases across the world. The extraction of the data is from the database [18].

### Mutation effect: Evolution of SARS-CoV-2 divergence

Mutations are considered as the basic units of species evolution [24], which are sometimes harmful and sometimes beneficial [26] but play important roles in genetic diversity via natural selection [25]. RNA viruses are generally mutation prone viruses during virus-host interaction, and their mutation rates are high (10^-4^ — 10^-6^) due to lack of proofreading activity, and the small increase in mutation rates in them may cause inability to evolve with the host [26]. It is also found that this high mutation rate intensifies virulence and coevolution of the virus with the host [24]. Further, these mutations in them are accountable for viral genome replication [27]. Hence high mutation rates in them have some significance, namely, in helping adaptability to the host [27], in enhancing the degree of viral genome replication kinetics [28] etc. leading to increase in their virulence causing human pandemic.

We have mined publicly available complete genomic data of various SARS-CoV-2 isolates (42 isolates in total) for seven countries (Supplementry TABLE 1). Taking Wuhan seafood market isolate with accession number NC_045512.2 as the reference genome, sequence comparison of these isolate genomes of SARS-CoV-2 with the reference genome is done, and clustered the isolates as phylogenetic tree (Fig. 2). The phylogenetic tree thus obtained (Fig. 2) indicates that there are approximately three main clusters of isolates (shades colors in Fig. 2) distributed across the countries. Surprisingly, it is also found that some of the isolates obtained in other countries (LR757996, MN988668, MN988669, MN996528, MN996530, MT019532) are completely similar to that of the reference genome. Further, the distribution of the virus isolates across these countries are found to be random.

**FIG. 2:**
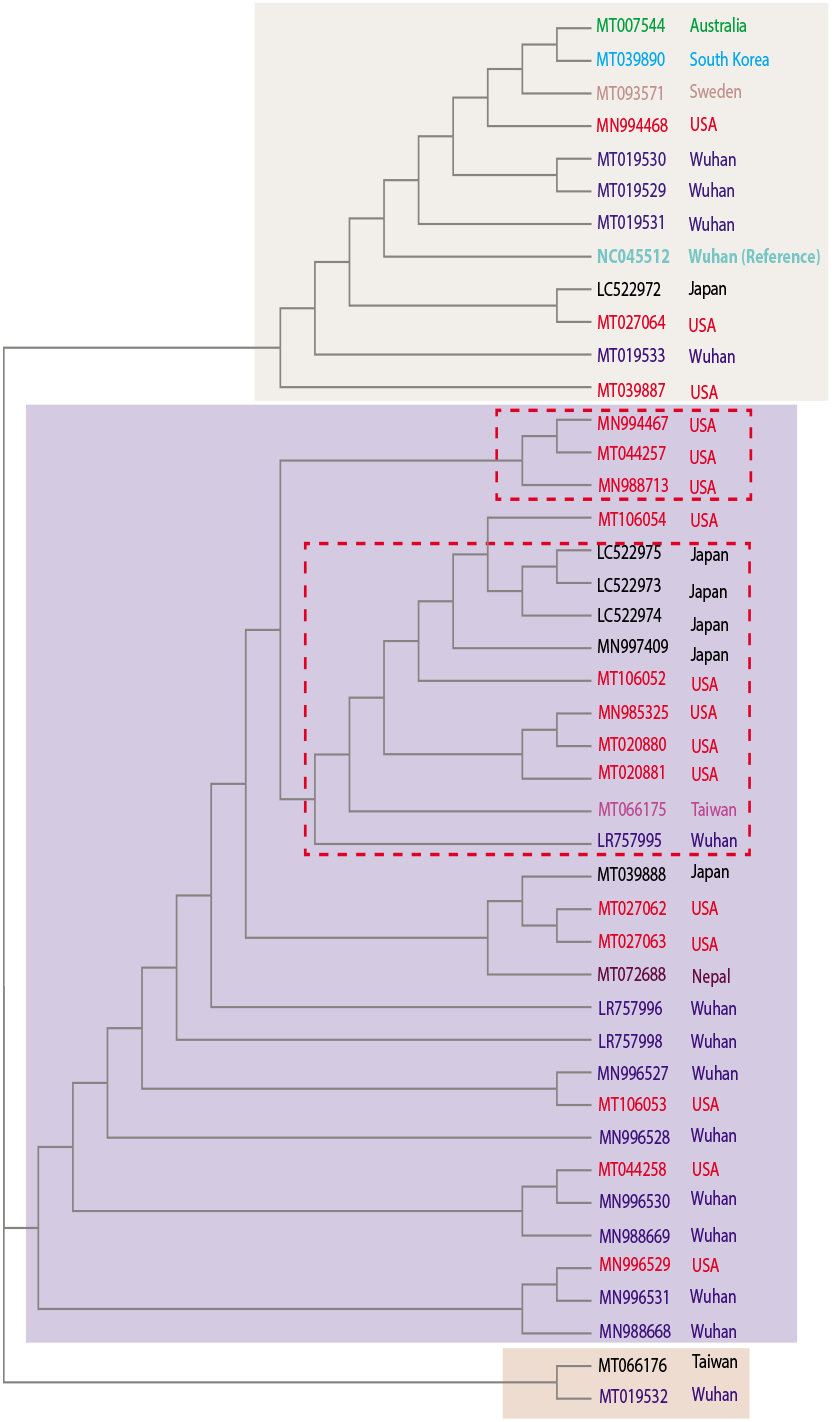
Phylogenetic tree of SARS-CoV-2 isolates: Clustering of SARS-CoV-2 virus isolates collected from the Covid-19 patients samples from eight different countries across the world.

We, then, calculated mutation rates (mutation per unit nucleotide) of the mined isolates, and found that the mutation rate (*m_r_*) of these isolates are very high (*m_r_* ∈ [10^-4^ — 10^-5^]), which could be a reason of causing the pandemic. The calculated mean value of the mutation per gene of the virus with error bars given by first and third quartiles (Fig. 2A) indicates that mutations take place at some specific genes only (mainly in *orf1ab, S, ORF3a, M, ORF8, N* genes). The results show that *m_r_* of *orf1ab* and *ORF3a*, such that *m_or f 1ab_*〉*m_ORP3a_*, and these two genes could be key genes for enhancing virus virulence and transmission [24, 27] which may lead to a disease epidemic. Further, the mean values of mutation rates of all isolates in each country are calculated and found the overall mutation rates S. Korea, Sweden, Japan, and the USA are large in comparison. The confirmed Covid-19 positive case, as well as death rate in these countries, are found to be comparatively large in comparison with other countries (Fig. 1), indicating an increase in mutation rate enhances the virulence of SARS-CoV-2 virus [24, 28]. Further, we calculated mutation rates of all six types of mutations *N_i_* → *N*_*i*-1_, where, *i* = 1,2, 3,4, *N_i_* ∈ [*A,C,T,G*] and, for the sake of simplicity, *N_i_* → *N*_*i*-1_ and *N*_*i*-1_ → *N_i_* are taken to be the same in the calculation (Fig. 3B). The results show that the average mutation rates of these mutation types in each country are found to be random and follow the Markov process [29]. It is observed that the transition is more frequent than transversion. During the evolution of genomes, the transition is more frequently observed as compared to transversion as the former usually does not lead to altering the protein, but the later has a higher impact on the change in the protein structure and function [30]. Some isolates of Wuhan, China contains transversion indicating that there is rapid evolution leading to adaptation with the new environment for survival or possibly making these isolates more virulent compared to their counterpart. On the other hand, *LR757995* out of a total of 15 isolates only consist of *C → T* mutation type at genomic location 8781. This specific mutation is seen to mutate along with mutation from *T → C* at genomic position 28143. These two mutations at positions 8781 and 28143 are highly prevalent in the USA (14 out of 16) and Japanese isolates (3 out of 4). Besides the isolates available from South Korea, Nepal, Sweden, one isolate from Taiwan and Australia does not contain these two specific mutations. It is probable that the genomic isolates spreading in the USA are less virulent compared to the reference Wuhan virus.

**FIG. 3:**
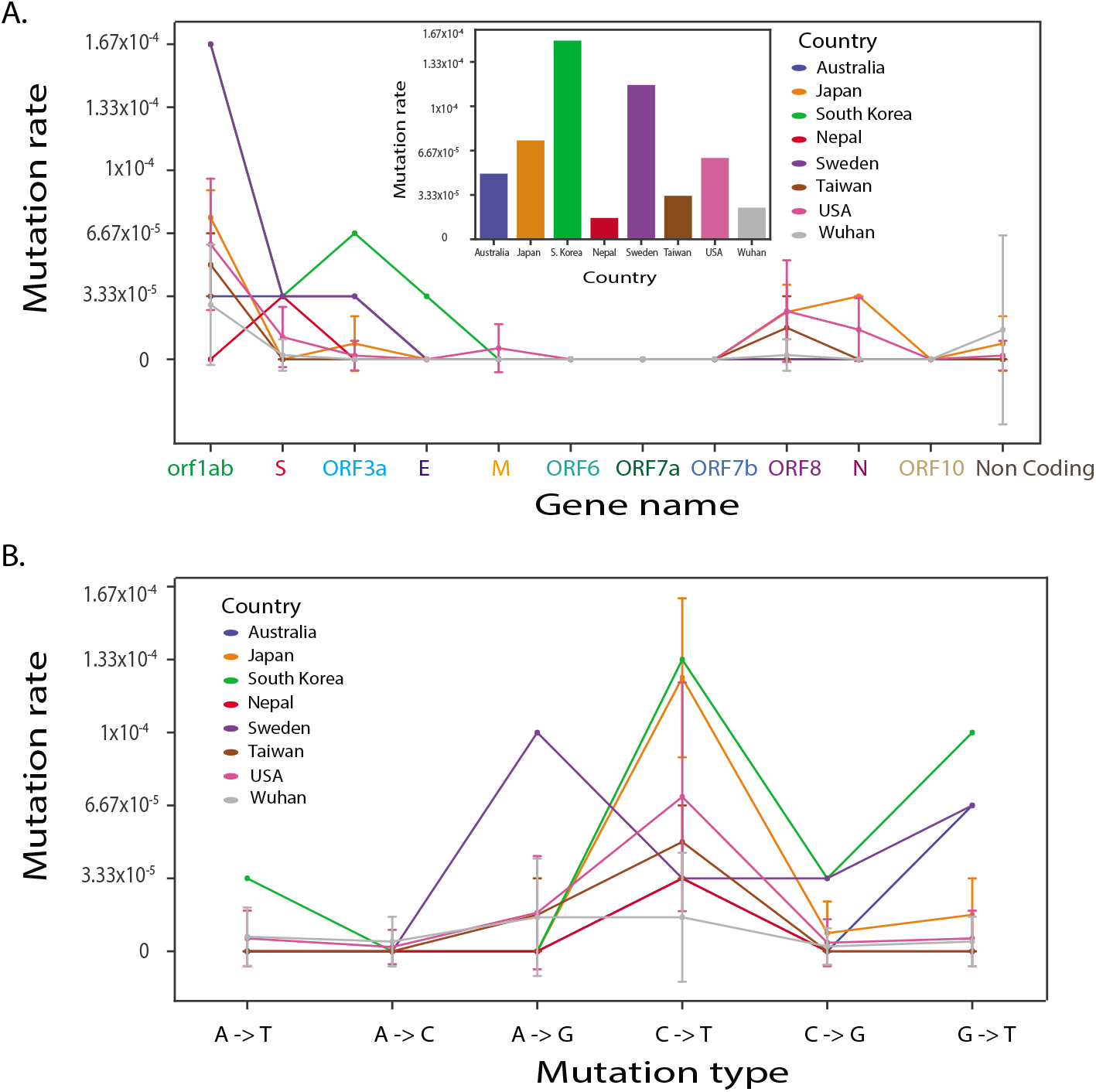
Mutation profiles of the SARS-CoV-2 isolates: (A) Plots of mutation rates verses gene name of SARS-CoV-2. (B) Plots of mutation rate with respect to mutation types.

On the other hand, out of all Japanese isolates available, only one isolate showed transversion mutation at genomic location 29704. This mutation is at the coding region and codes for the 3’ stem-loop II-like motif (also known as s2m) whose function is not yet fully known, but have been found to have a role in viral replication [31]. But we observed that most of these isolates have transitional mutation *C → T* at genomic location 8781. This mutation falls in the coding region, namely, *nsp4* and codes for transmembrane domain 2(TM2). Maximum mutation type is observed to be *C → T* in most genomic isolates falling in the orf 1ab gene (Fig. 3A and B).

### Complexity in SARS-CoV-2 isolates

We analyze the genomic complexity of the SARS-CoV-2 isolates using multifractal analysis [32, 33] to get an insight into the impact of genome variability and to understand the virulence of the virus isolates in a different perspective. In this technique, we first convert the symbolic genome sequence to time series like DNA walk by taking purine (A or G) as step-up (+1) and pyrimidine as step down (− 1) [see the *Methods*]. DNA walk is a cumulative sum, which means that the effect of a single point mutation can lead to different walk altogether, and hence we can quantify them using Multifractal parameters [see the *Methods*]. Then multifractal analysis can be done from the DNA walk of each viral isolate to calculate various multifractal parameters. The *DNA walks* of all SARS-CoV-2 RNA virus isolates collected from the patients’ data from various countries (42 isolates) as a function of order parameter q obey Markov process (Fig. 4A). The calculated *fluctuation function F_q_*(*s*) of order *q* as a function of normalized *s* [*s* = *int*(*S/L*), *L* is the length of the DNA walk of the virus genome] for all SARS-CoV-2 virus isolates follow overall power-law behavior *F_q_*(*s*) ~ *s^H_q_^*, where, *H_q_* is the Hurst exponent of order q [34] (Fig. 4B) indicating fractal nature in the virus genome organization. Further, the calculated *H_q_* as a function of *q* is found to be *q* and *s* dependent (Fig. 4B and Fig. 5D) indicating the scaling nature in *F_q_*(*s*) is inter-event dependent showing multifractal nature [32]. Since the *F_q_* behavior indicate *local* multiple fractal natures at various local domains of s with various local Hurst exponents, 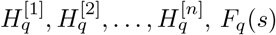 exhibits self-affine multifractal property via observable variable *z*(*s*) [32, 33, 35] given by,

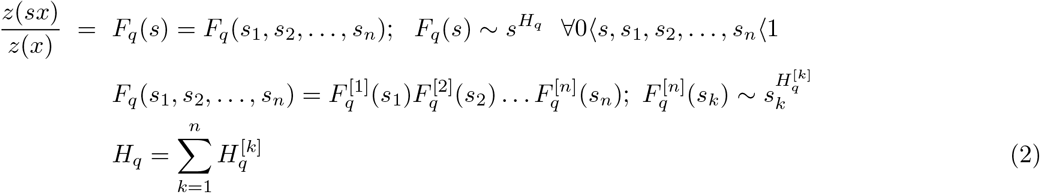

*not clear* And each local segment *s_k_* satisfy fractal law [33]. We then calculated the average 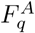 of all virus isolates in each country with error bars indicated by first and third quartiles (Fig. 4C). We found that 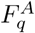 in S. Korea is much higher than in other countries, and 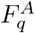 variability is much more in Wuhan and then in Japan. The high value of 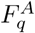 indicates the presence of multiple isolates in Wuhan, and induces large variability in the virus genome could cause large variability in 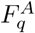 which might be harm to the hosts in the disease epidemic.

**FIG. 4:**
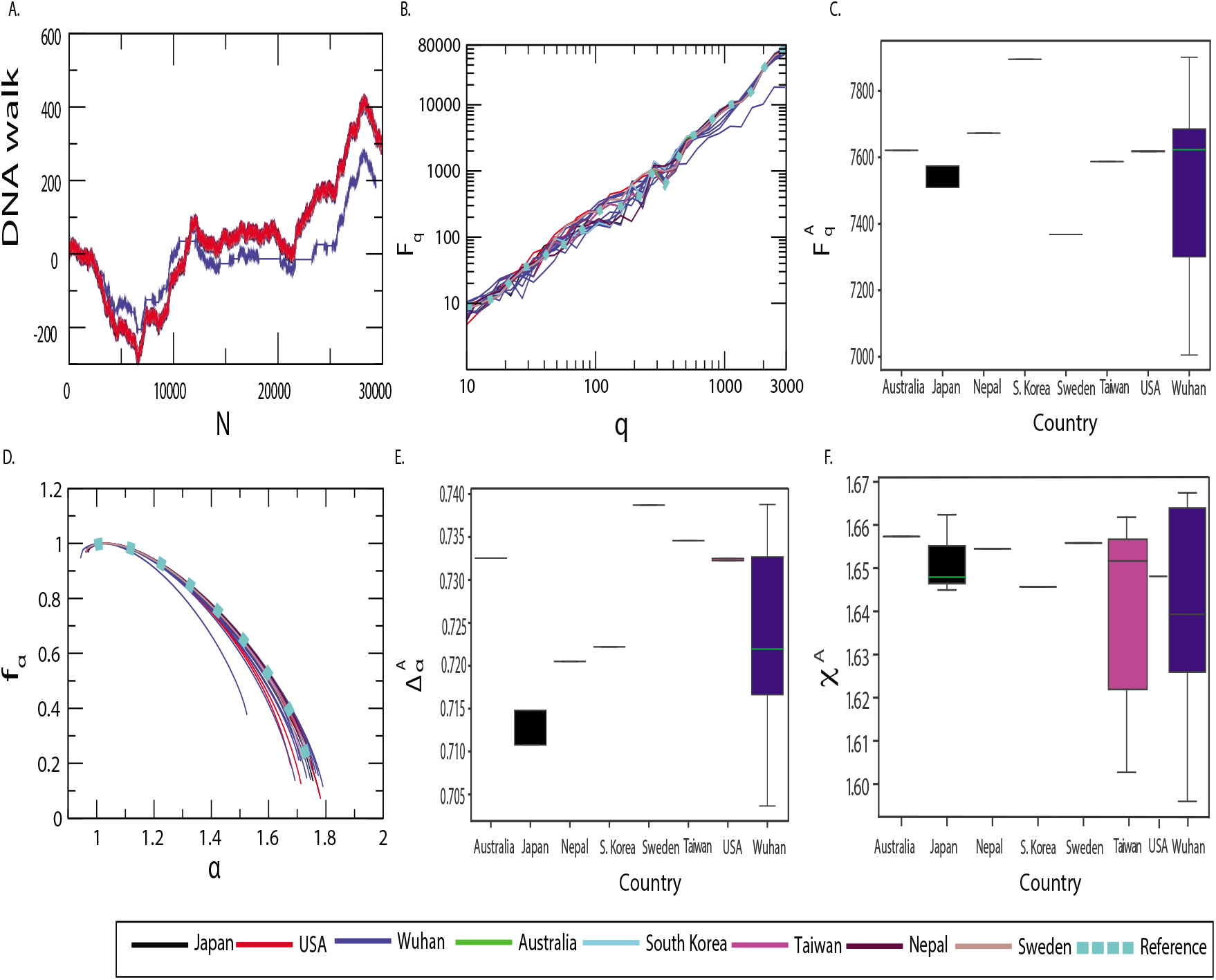
Multifractal parameters of the SARS-CoV-2 isolates: Plots of (A) DNA walks of all the 43 genomic isolates of SARS-CoV-2 where reference genome is shown as dotted light blue. (B) *F_q_* verses *q* in log-log plot, (C) averaged *F_q_* with respect to respective countries across the world, (D) *f_α_* as a function of *a* with dotted line being the reference, (E) Δ*α* for various countries, and, (F) averaged *χ^A^* for different countries.

We then calculated singularity function *f* (*α*) of all the virus isolates (Fig. 4D) and found the curves collapse roughly at *α* ~ 1 and left and right hand parts of of the curves around the value of *α* = *α*\* at which *f* (*α*\*) → *f_max_* (*f_max_* is the maximum value of *f*) is quite biased and even left parts of the curves about *α*\* are quite small. Since *f* (*α*\*) → *f_max_* for *α*\* ~ 1.13 ± 0.012)1 for all curves, the virus isolates are of heterogeneous structure driving genome organization complex [36] (Fig. 4D). To understand the behavior of f around *α*\*, we can expand *f* (*α*) around *α*\* by Taylor series expansion,

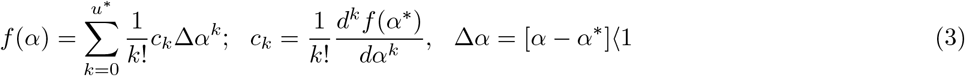

where, *u*\* is the degree of *f* (*α*) after truncation. Then the symmetry of *f* (*α*), which can characterize the complexity in the system, is defined by a skew parameter *χ*,

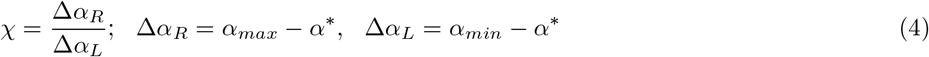

where, *f* (*α*) is right-skewed if *χ*〉1, left-skewed if *χ*〈1 and perfect symmetry if *χ* = 1 [32]. It is clearly seen from Fig. 4D that Δ*α_R_* ≫ Δ*α_L_* indicating asymmetric nature of *χ*. Hence, we calculated average 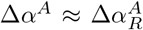 of each country taking into account all the virus isolates in the country (Fig. 4E), and found that the value of Δ*α^A^* is much higher in Sweden, Australia, Taiwan and USA indicating richness in genome structure showing strong signature of multifractality, leading to the complex behavior of the virus. However, the variability of Δ*α^A^* is quite large in Wuhan, which could be due to the presence of multiple isolates indicating virulence of the virus leading to epidemic [37, 38]. Then we calculated average *χ_A_* for the countries we have taken and found that *χ^A^*〈1 (left-skewed) revealing the dominance of the virus genome organization and scaling in multifractality by large fluctuations [32, 36] (Fig. 4F). The values of *χ^A^* are found to similar for all the countries, but the variability is much large for Wuhan, Taiwan and Japan which could be dependent of the large fluctuations in the virus isolates on demography.

Now, we calculated generalized dimension *D_q_* for the virus isolates from their corresponding DNA walks using,

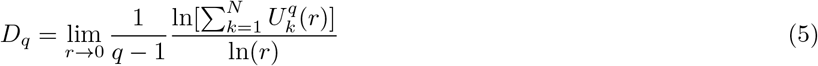

where, 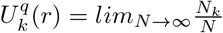 is the probability that kth DNA walk segment of length scale *r* will have *N_k_* observations, and obeys 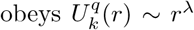 [34], where, λ is Holder exponent [39]. The results of *D_q_* shows that *D_q_* is quite dependent on *q* (Fig. 5A) indicating rich heterogeneous structure in the virus isolates [32], where, local structures in the genome could have various properties and functions. The changes in those local structure may induce drastic change in genome as well as phenotypic behavior of the virus isolates. We then calculated relative change of the isolates with respect to reference virus genome 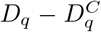 with *q* (Fig. 5B) shows significant changes in the parameters and 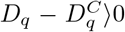 indicating increase in complexity due to increase in heterogeneous structure as compared to reference virus genome. The complexity in virus isolate in Japan is quite high (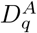 is highest) as compared to other countries (Fig. 5C), and variability in complexity in Wuhan is much more as compared to other countries.

**FIG. 5:**
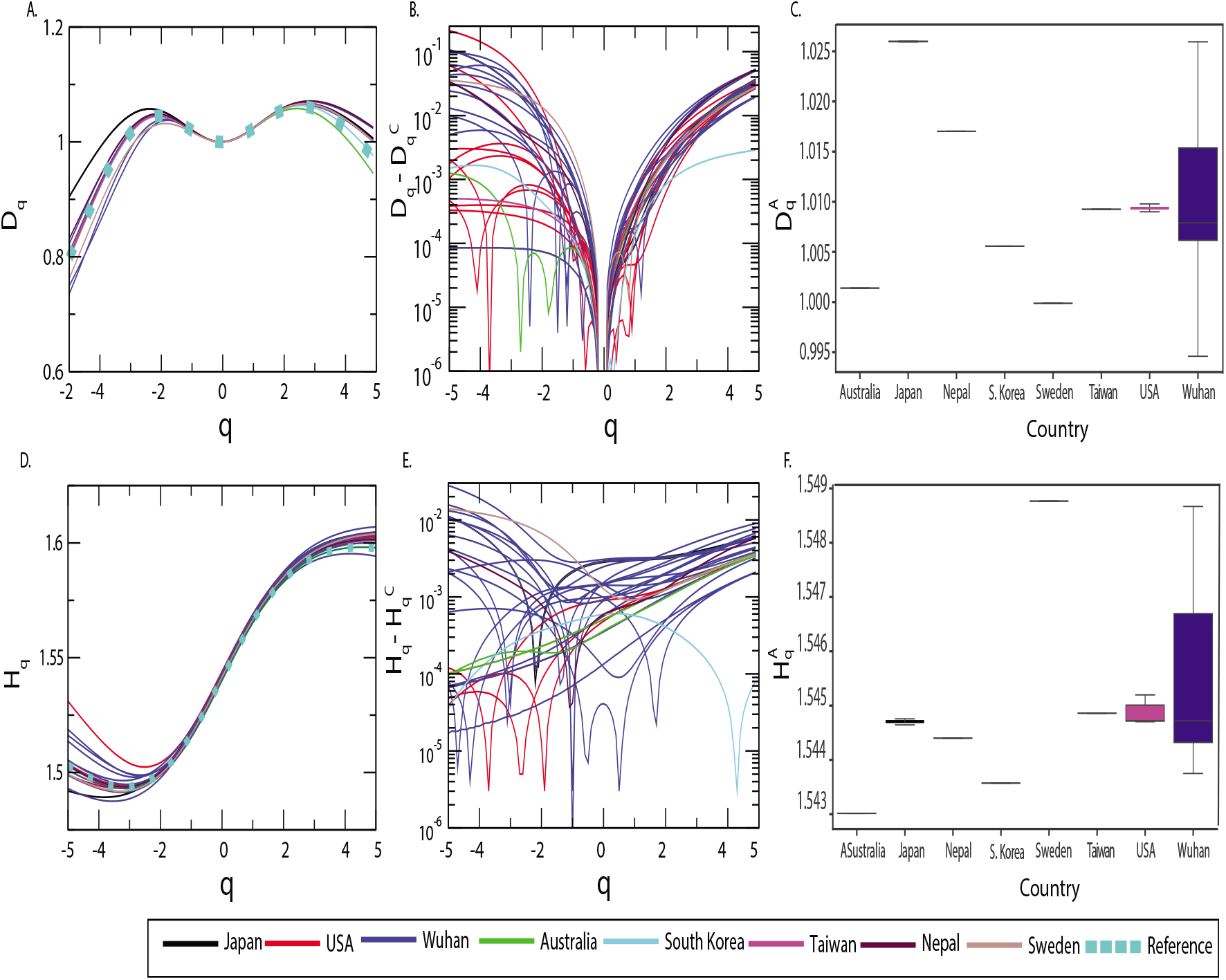
Nature of *D_q_* and *H_q_* of the SARS-CoV-2 isolates: Plots of (A) *D_q_* versus *q* in log-log plot for 43 genomic isolates of SARS-CoV-2 where reference genome is shown as dotted light blue, (B) averaged *D_q_* versus *q* in log log plot, (C) averaged *D_q_* as a function of respective country, (D) *H_q_* with respect to *q* with dotted line being the reference, (E) 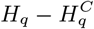 versus *q*, and, (F) averaged *H_q_* for the countries we have taken.

We, then, calculated *H_q_* as a function of *q* (Fig. 5D) by using the MFDFA approach described in *Methods.* Since the *H_q_* values are quite variable with respect to *q*, the isolate genomes show high heterogeneous structures. Further, since *H_q_*〉0.5, genomic signal in the whole virus isolate genome sequences indicate positive correlations in them [40] indicating possibility of strong self-organizing behavior in each virus isolate [32, 41]. The calculated change in *H_q_* with respect to reference virus genome 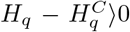 is positive (Fig. 5E), the correlations in the virus genome increases in the mutated isolates which could be one reason for increase in virulence in virus isolates. The average value in 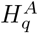 and variability are much more in Wuhan virus isolates as compared to other countries (Fig. 5F). The multifractal analysis of the virus isolates across the countries reveals that mutated virus isolates have increased virulence as compared to the reference virus. Further, presence of multiple isolates in a host also enhances virulence of the virus. The virus isolate is quite sensitive to fluctuations and might correlated to increase in virus mutations which cause heterogeneity in genome structure exhibiting strong multifractal properties.

### Price theory of evolution of mutant isolates in infected disease

One of most important basic reasons of outbreak of an epidemic could be due to sudden trade-off of multiple viral species modulated by high mutation effects, which obey Markov process, due to virus-hosts interaction [26, 42]. This epidemic process could be dependent on various factors, namely, virus virulence, transmission (passive and active) etc [43]. Let us consider *M* mutationally different isolates of the virus in the hosts, and *x_i_* be the quantitative trait of ith individual virus or parasite (0 ≤ *x_i_* ≤ 1) in the hosts, then the dynamics of mean of this variable 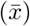 is given by Price equation [43–45],

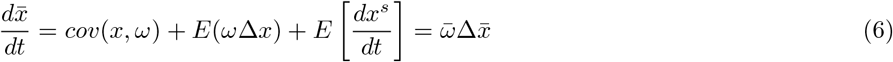

where, 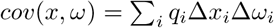 and 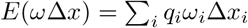 are the covariance and expectation value functions, with *q_i_* as the frequency and *x^s^* as the expected value of survived individual virus during [*t,t + dt*] respectively [44]. The model we consider incorporates the mechanism of infecting susceptible hosts S by a virus isolate of type ‘ *i*’ (*y_i_* is the concentration of the ith isolate) and then turn into new dominating mutant isolate type ‘*j = i* + 1’ due to mutation event in hosts-virus interaction [46]. The model allows the presence of different isolate types during the interaction with the host causing more virulent harming the hosts which could be one of the most important factors which could drive the virus transmission among the hosts [37, 38]. Then the dynamics of ith isolate variable *y_i_* is given by,

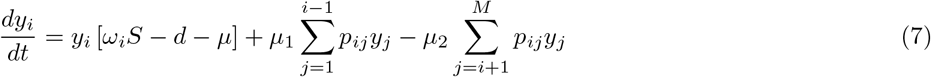

where, *i* = 1, 2, …, *M*, *ω_i_* corresponds to fitness parameter, which is the transmission rate at which hosts infected by isolate ‘i’ infect *S, d* is the death rate of hosts, *μ*_1_ is the rate of generating self mutant ‘i’, *μ*_2_ is the rate of generating new dominating mutant isolate ‘*j*’ and *p_ij_* is the probability that infectious isolate ‘ i’ turned into isolate ‘j’ and *μ* = *μ*_2_ – *μ*_1_. Then we can define, 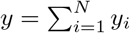, and frequency, 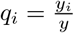, and equation (7) becomes, 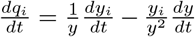. Now substituting (7) to this expression, using 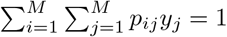 and after simplification, we arrived at the following equation,

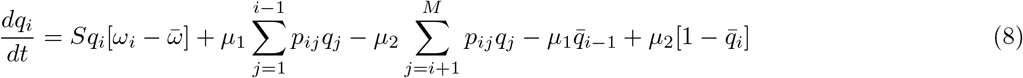

where, 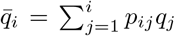. Now, the mean 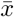 can be obtained from 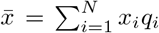. Then using to the equation (8), simplified the obtained equation and rearranging the terms, we have the dynamics of 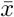 of the model by the following Price equation,

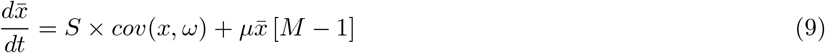

The fitness parameter of individual virus isolate type i of jth parent is composed of two forces arises from competition in the host and increase virulence, given by, 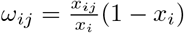 [43]. Hence, the fitness of the ith mutant isolate group will be given by, *ω_i_* = (1 – *x_i_*). Now, this *ω_ij_* become one important parameter that determine virus isolate trade-off which can in general represented by, *ω* = *xf* (*x*) [42], such that, 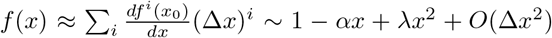, where, 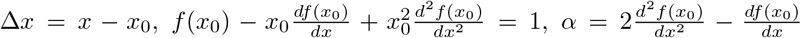 and 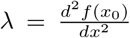. Then, one can approximate fitness parameter to be, *ω* ~ *x* [1 — *αx* + λ*x*^2^]. The optimized fitness of the virus can be obtained by calculating the equilibrium state of *ω* and is dependent on various factors to control the virus transmission/diffusion [42, 47].

Now, if one consider *κ_i_* is the genetic variability (additive in nature) of the ith mutant isolate group, then, *x_i_* can be expressed as, *x_i_* = *x*+*∊κ_i_*, where, *∊*〈1 [43]. Now, we have, Δ*x* = *x_i_*—*x* = *∊κ_i_*, and Δ*ω* = *ω_i_*—*ω* = (1–*x_i_*) –(1—*x*) = —*∊κ_i_*. Substituting these values, we have, 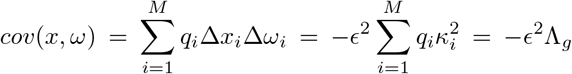, where, Λ_*g*_ is the genetic variance of the mutant isolates in the group. Now, putting back to the equation (12) and rearranging the terms, we have,

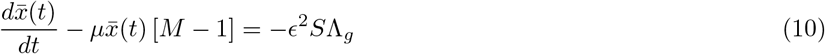

This is special Bernoulli differential equation with constant coefficient and inhomogeneous term, which can be solved easily by the technique of calculating integrating factor (*IF*), where, *IF* = *e^-μ(M-1)t^*, or by simple separation of variables method. Then after solving the equation (10), the expression for 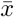 is given by,

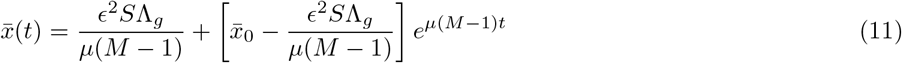

where, 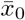 is the amount of 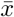 at *t* = 0.

#### Theorem 1

*Optimal trait value has the properties:* 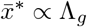 *and* 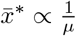.

**Proof:** *The optimal trait value* 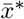 *can be obtained by taking* 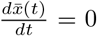 *in equation* (10), *or by taking* lim_*t*→0_ *x*(*t*) *in equation (11). It is given by*,

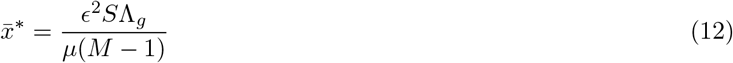

*From this equation (12), we can directly obtain*, 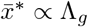, and 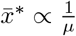.

#### Lemma 1

*Phase plane analysis of the virus-pathogen interaction system modeled by* (10) *shows that mutational rate drives the system an “Asymptotic stability”.*

**Proof:** *The equation (10) can be written as* 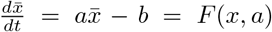, *where, a = μ* [*M* — 1] *and b* = *∊*^2^*S*Λ_*g*_. *Fixed point of the dynamical system is given by*, 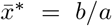, *and hence*, 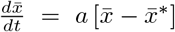. *The stability of the dynamical system around a fixed point can be analyzed by calculating*, 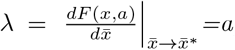, *such that, for* λ〉0, 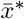 *is unstable, whereas, for* 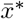 *is stable [49]. Phase plane analysis of this dynamical system can be done as follows: (a) for a〉0: (i)* 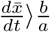 *if* 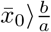, *such that* 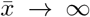 *departing from* 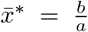; (*ii*) *for* 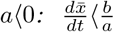 *when* 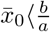, indicating 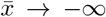; *and (iii) for* 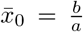 *shows* 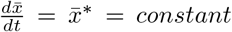. *Therefore, the system is unstable, and the system will diverge for slight perturbation to it. Further*, 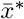, *in this case, is unstable fixed point because* λ〉0 *for a〉0. This condition corresponds to the Malthusian explosion. Following a similar argument for the case a〈0, it can be shown that* 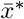 *is a stable fixed point. Hence, the bifurcation diagram of this dynamical system exhibit asymptotic stability (Fig. 6). Since*, *μ* = *μ*_2_ – *μ_1_*, *we have, a* = (*μ*_2_ — *μ*_1_) [*M* – 1]. *The bifurcation diagram indicates that for μ_2_*〉*μ_i_*, 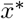 *is unstable fixed point. This is the condition where the rate of generating new dominating mutant isolates is much more than that of generating its own isolates. This condition could lead to the trade-off of the new mutant isolates away from the original hosts. This could be the origin of the spreading of viral infection due to the generation of new mutant viral isolates. This claim can be correlated to the notion that since mutations in RNA virus is high [26, 27] causing harm to the hosts and hence, transmitted by diffusing from one host to another causing infection to other hosts [28]. Otherwise, the old viral isolates will stick to the old hosts* 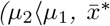 *is a stable fixed point since λ〈0) without diffusing from the hosts. Further, the characteristics time scale, which is the time required to determine significant change in* 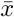 *around fixed point* 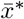, *is given by*, 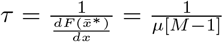.

**FIG. 6:**
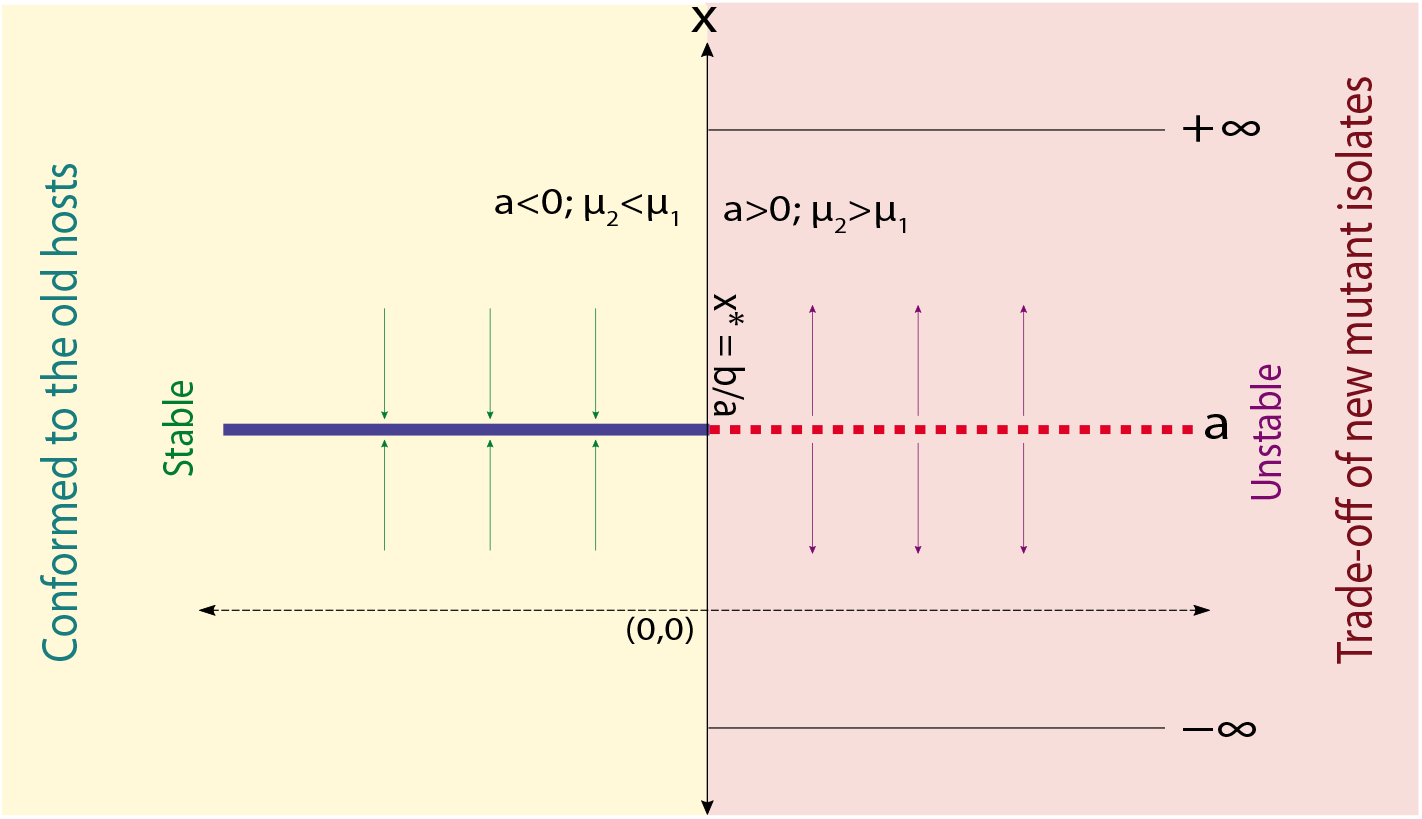
Bifurcation diagram of the virus-hosts interaction model.

#### Lemma 2

*Competition might drive the migration of virus isolates to different hosts.*

**Proof:** *The criteria of competitive or cooperative of any two mutant isolates modeled by the dynamical system*, 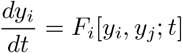, *is given by [50]: 1. The mutant isolates ‘i’ and ‘j’ are competitive in a region R if*, 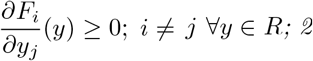. *The two mutant isolates are cooperative if*, 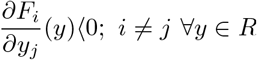. *Applying this criteria to the model we considered* (7), 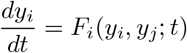, *we have*,

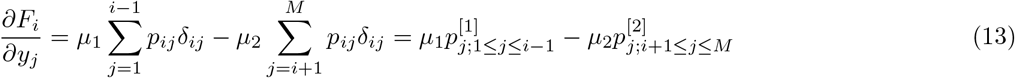

*such that, ‘i’ and ‘j’ mutant isolates competitive if* 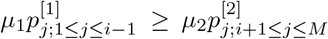 *This condition allows the two mutant isolates compete among themselves, and if the new mutant ‘j’ dominates over ‘i’, then there could be high chance of more virulence due to it’ s mutation and may trade-off from the present hosts to the others. However, if* 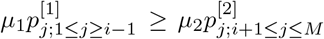, *then the two mutant isolates will cooperate to each other and may confine to the present hosts. This condition could happen when the virulence of the two mutant isolate types are nearly similar due to not much difference in the mutation profiles.*

## Conclusion

SARS-CoV-2 is a highly virulent RNA virus causing pandemic across the world. Some of the most important factors of the origin of this virulence could be due to high mutational events in the virus genome when hosts-virus interaction takes place, and the presence of multiple virus isolates in the host. The Covid-19 pandemic is all of a sudden first epicenter from Wuhan, China, and spread all over the work, killing nearly two lakh people across the world. The pandemic is so fast, no concrete theory of the widespread of this disease, and still no vaccine is discovered yet. In this work, we studied and analyzed a large number of publicly available SARS-CoV-2 genomes collected from patients from different parts of the world to explore the possibility of the origin of this pandemic. Our study of mutational profiles of SARS-CoV-2 isolates across the world show very high mutational rates in these isolates, which led the isolates more virulent, causing significant harm to the hosts and enhance viral transmission to hosts. The mutations in these isolates also take place mainly in six specific genes out of eleven genes, and the highest mutation takes place in the *orf1ab* gene and the mutation events follow the Markov process. It is also found from the mutational profile study that transition is much more frequent than transversion.

We, then, analyzed the SARS-CoV-2 isolates using the multifractal approach to understand the complexity in the isolates genome variability and structural organization. The calculated multifractal parameters reveal that structural variability in the isolates is caused by the presence of multiple isolates in the hosts. These isolates are highly asymmetric (left-skewed) in the singularity function, indicating the richness of complexity and dominance by large fluctuations in genome structure organization imparting virulence in the virus. The large values obtained for Hurst exponent and generalized dimension indicates heterogeneous genome structure organization and strong positive correlation in the organization of SARS-CoV-2 isolates. The genome structures of the isolates are far from equilibrium, self-organized, and quite sensitive to fluctuations in and around it. This could also be the origin of multiple isolates in a certain geographical area. The presence of multiple isolates in hosts may lead the Covid-19 pandemic.

We then study a simple model of hosts-virus interaction by taking into consideration multiple isolates in it and applied Price theory to understand the epidemic. The theory indicates that the fitness parameter of the isolates, which is dependent on various factors, namely virulence, fluctuations, etc., is quite sensitive and affected by fluctuations, multiple isolate interactions with the host, etc. However, Price theory needs to be investigated by taking into account various factors such as quarantine, exposed, health care, optimized economy, and many other factors.

## Materials and Methods

The detailed methods of this study are given below.

### Conversion of genomic sequence into DNA Walk

The whole genome of every isolate as well as the reference genome are converted in *DNA walk* {*x_i_; i* = 1, 2,…, *N*} by considering purine (A or G) as step up (*x_i_* =+1) and pyrimidine (C or T) as step down (*x_i_* =-1) [48]. The *DNA walk* is considered to be a non stationary time series data due to its stochastic behavior and is the input considered for various parameters. This DNA walk is considered as the profile, 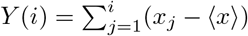, where, 〈*x*〉 is the mean, and *i* = 1,2,…,*N*.

### Multifractal Detrended Fluctuation Analysis (MF-DFA)

The Multifractal Detrended Fluctuation Analysis (MF-DFA) is a powerful method for analysing and studying fractal characteristics in non-stationary time series data[34]. Using the numerical technique mentioned in Kantelhardt et al.[32, 33] various multifractal parameters that describe the fractal nature in the time series can be calculated such as generalized dimension(D_q_), Hurst exponent(H_q_),singularity spectrum(*f* (*α*)) etc. We calculate these parameters for all the three types of isolates. The procedure of MF-DFA has five steps which are summarized.

- Step 1. We consider the nonstationary time series data as the input to MF-DFA algorithm for calculating all the multifractal parameters. In the first step in MF-DFA the DNA walk time series signal *x_j_* having length *N* is considered and profile representing the time series is calculated as below,

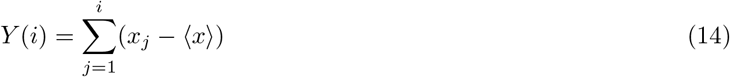

where < *x* > is the average value and i = 1,2,3,…N.
- step 2. The profile *Y*(*i*) is divided in non-overlapping segments of length (l) of same size as *N_s_* ≡ *int*(*N/s*). It is to note that as the length N of the time series is not usually the multiple of examined time scale s therefore at the tail part of profile some small part could be remaining. Keeping this in mind the same protocol is redone from the other end of the time series. Therefore 2*N_s_* segments are calculated altogether.
- step 3. Using the least-square method of fitting calculate the local trend for every segment(2*N_s_*). We calculate the variance as follows,

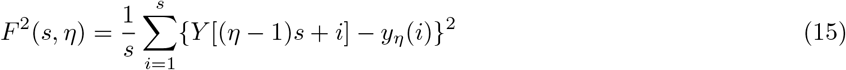

where for every segment *η, η* =1, 2, 3,…, *N_s_* and

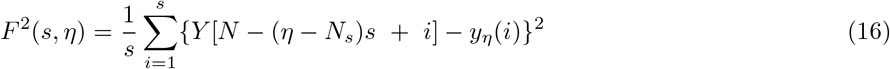

for *η* = *N_s_* + 1,…, 2*N_s_* and *y_η_* (*i*) is the polynomial that will be fitted in the segment *η*.
- step 4. The fluctuation function of *q^th^* order is estimated using the average of all the segments as,

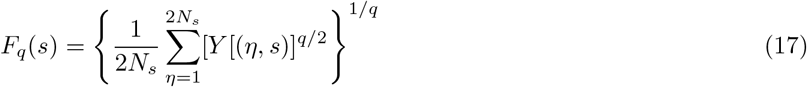

where, *q* an index variable is any real valued number except zero[33].
- step 5. The scaling behaviour of fluctuation *F_q_*(*s*) is calculated after analysing the log-log plots (*F_q_*(*s*) with respect to *s*) for a single *q* value and if time-series is has long range power-law correlation then *F_q_*(*s*) will increase for larger values of *s* following power law as,

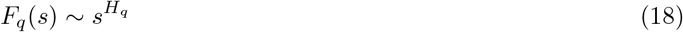

The *H*(*q*) is the generalised hurst exponent and in our analysis we have calculated the *H*(*q*) by considering *q* from *q_max_* = —5 to *q_max_* = +5. The value of *H*(*q*) changing with the change in value of *q* represents that our DNA walk data multifractal nature and it is to note that if varying the value of *H*(*q*) with respect to *q* is constant then it indicates that time series is monofractal in nature.

The relationship between *H*(*q*) and mass exponent is calculated for the various isolates as follows,

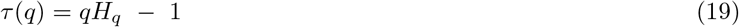

Another important multifractal parameter is the singularity spectrum for characterizing a multifractal time series. It is related to *τ*(*q*) with the Legendre transformation. *α* and *f* (*α*) are calculated as below,

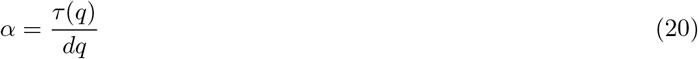

and

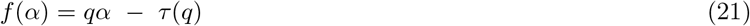

Here, *α* is called the Holder exponent also called singularity strength and *f* (*α*) is the singularity spectrum and can be simply expressed in the form of *q* and *H*(*q*) as the following equation[33].

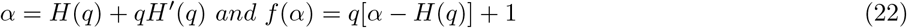

The generalised multifractal dimension (*D_q_*) is another multifractal parameter and is defined as follows[33],

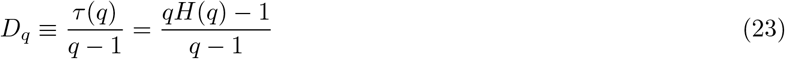

here, *τ*(*q*) is Mass exponent and *H*(*q*) is Hurst exponent. where, The *f* (*α*) is the subset dimension of the series characterised by *α* whereas the *α* is called Holder exponent or singularity strength. We performed the alignment of all the genomes and find out the point mutation present in all the isolates with respect to the reference Wuhan genome and calculated the various multifractal parameters.

## Acknowledgement

SM (Project ID: 2019-5116, File No: ISRM/11(16)/2019) is financially supported by Indian Council of Medical Research(ICMR). MZM financially supported by Department of Health and Research, Ministry of Health and Family Welfare, Government of India under young scientist FTS No. 3146887. R.K.B.S. acknowledges UPE-II, sanction no. 101, India, for providing financial support.

## Author Contributions

RKB and MZM conceptualized and designed the model. SM, RKSS, SKS, MZM and RKBS did the computational experiment and prepared the figures from the results. SM, RKSS, SKS, MZM and RKBS wrote the manuscript. All authors read, checked and approved the manuscript.

