## Supplemental Table 1 for "Complexity in SARS-CoV-2 genome data: Price theory of mutant isolates": Supplementry.pdf

TABLE I: Genomic Information

| S.No | Accession No. | Region | No. of Transition | No. of Transversion | Total Mutations |
|---|---|---|---|---|---|
| 1 | LC522972 | Japan | 2 | 2 | 4 |
| 2 | LC522973 | Japan | 5 | 0 | 5 |
| 3 | LC522974 | Japan | 4 | 0 | 4 |
| 4 | LC522975 | Japan | 4 | 1 | 5 |
| 5 | LR757995 | Wuhan | 3 | 0 | 3 |
| 6 | LR757996 | Wuhan | 0 | 0 | 0 |
| 7 | LR757998 | Wuhan | 0 | 2 | 2 |
| 8 | MN985325 | USA | 3 | 0 | 3 |
| 9 | MN988668 | Wuhan | 0 | 0 | 0 |
| 10 | MN988669 | Wuhan | 0 | 0 | 0 |
| 11 | MN988713 | USA | 7 | 0 | 8 |
| 12 | MN994467 | USA | 5 | 2 | 7 |
| 13 | MN994468 | USA | 1 | 1 | 2 |
| 14 | MN996527 | Wuhan | 2 | 0 | 2 |
| 15 | MN996528 | Wuhan | 0 | 0 | 0 |
| 16 | MN996529 | Wuhan | 2 | 0 | 2 |
| 17 | MN996530 | Wuhan | 0 | 0 | 0 |
| 18 | MN996531 | Wuhan | 1 | 1 | 2 |
| 19 | MN997409 | USA | 3 | 1 | 4 |
| 20 | MT007544 | Australia | 1 | 2 | 3 |
| 21 | MT019529 | Wuhan | 2 | 1 | 3 |
| 22 | MT019530 | Wuhan | 3 | 3 | 6 |
| 23 | MT019531 | Wuhan | 1 | 0 | 1 |
| 24 | MT019532 | Wuhan | 0 | 0 | 0 |
| 25 | MT019533 | Wuhan | 0 | 1 | 1 |
| 26 | MT020880 | USA | 3 | 0 | 3 |
| 27 | MT020881 | USA | 3 | 0 | 3 |
| 28 | MT027062 | USA | 3 | 0 | 3 |
| 29 | MT027063 | USA | 3 | 0 | 3 |
| 30 | MT027064 | USA | 2 | 0 | 2 |
| 31 | MT039887 | USA | 1 | 0 | 1 |
| 32 | MT039888 | USA | 4 | 1 | 5 |
| 33 | MT039890 | South Korea | 4 | 5 | 9 |
| 34 | MT044257 | USA | 5 | 2 | 7 |
| 35 | MT044258 | USA | 0 | 0 | 0 |
| 36 | MT066175 | Taiwan | 2 | 0 | 2 |
| 37 | MT066176 | Taiwan | 2 | 0 | 2 |
| 38 | MT072688 | Nepal | 1 | 0 | 1 |
| 39 | MT093571 | Sweden | 4 | 3 | 7 |
| 40 | MT106052 | USA | 4 | 0 | 4 |
| 41 | MT106053 | USA | 1 | 0 | 1 |
| 42 | MT106054 | USA | 5 | 2 | 7 |
